# Gain Control of Peripheral Input by Core Brain Thermosensors Drives *Drosophila* Thermoregulation

**DOI:** 10.64898/2026.11.03.764052

**Authors:** Stanislav Lazopulo, Marta Zlatic, Albert Cardona, Paul A. Garrity, Aravinthan D.T. Samuel

**Affiliations:** Department of Physics and Center for Brain Science, Harvard University, Cambridge, MA, USA; University of Cambridge, Department of Zoology and MRC Laboratory of Molecular Biology, Neurobiology Division, Cambridge, UK; Department of Biology and Volen National Center for Complex Systems, Brandeis University, Waltham, MA, USA

## Abstract

Temperature affects all aspects of animal physiology, rendering thermoregulation critical for survival. In *Drosophila* larvae, well-established peripheral thermosensors detect temperature changes and mediate thermotaxis over a broad innocuous range, but how larvae seek a specific preferred body temperature within this range remains unknown. Here we show that core thermosensors in the brain (LPBT neurons) establish this setpoint by mediating warming-evoked turning at temperatures above it. LPBTs express the warmth-activated thermosensor TrpA1(A), but are not simply cell-autonomous thermosensors. Rather, sustained LPBT responses above setpoint require both peripheral thermosensory input and TRPA1-dependent thermosensing within LPBTs. TRPA1-dependent core thermosensing provides gain control that defines the homeostatic setpoint by tuning the behavioral weights of peripheral inputs. This new conception of thermal preference reveals parallels between Drosophila and mammalian thermoregulation.

## Main Text

The maintenance of appropriate body temperatures (thermoregulation) is critical for animal survival. Homeotherms thermoregulate using both autonomic and behavioral mechanisms, including blood flow regulation, shivering and movement(*1–3*). Poikilotherms like *Drosophila* primarily rely on behavioral thermoregulatory strategies, such as locomotion toward favorable thermal environments (thermotaxis) (*4*). In both cases, to mount an appropriate thermoregulatory response, the animal must integrate multiple thermosensory signals, reflecting the temperature of the environment and of the animal’s own body. How and where this integration occurs is a fundamental question in body temperature homeostasis.

In mammals, temperature is sensed through multiple classes of thermosensory neurons, peripheral and central, that respond to diverse thermosensory stimuli ranging from innocuous warmth and cooling to noxious heat and cold (*5*, *6*). This perceptual range is spanned by molecularly distinct sensors with distinct thermosensitivities, including TRP-family cation channels(*7–13*), TREK two-pore potassium channels and the kainate-type ionotropic glutamate receptor GluK2(*14*, *15*). The distributed thermal signals from these diverse sensors must be integrated to generate coherent thermoregulatory responses. The preoptic area (POA) of the hypothalamus is a central node that receives ascending thermal information from peripheral sensors to coordinate both autonomic and behavioral responses(*1*, *5*, *16*, *17*). The POA also contains intrinsically temperature-sensitive neurons that are functionally distinct from rapid peripheral sensors of temperature change(*16*, *17*). Such internal sensors are well-suited to report absolute body temperature, as their location renders them relatively insensitive to rapid environmental fluctuations(*2*). Peripheral sensors, by contrast, are well-suited to rapidly respond to external temperature changes. Despite ongoing progress in identifying thermosensors, an understanding of how internal and peripheral thermosensory inputs are integrated remains elusive.

*Drosophila melanogaster* offers a tractable system for dissecting thermosensory integration. Like mammals, flies use distinct cellular and molecular sensors for different temperature regimes(*18*). For example, multiple TRPA-family channels contribute to noxious heat responses(*19–22*), while PKD2, TRPM, and NOMPC contribute to noxious cold responses(*23*, *24*). Away from the extremes of noxious avoidance, *Drosophila* larvae navigate toward a specific preferred temperature using dorsal organ cooling cells (DOCCs) and dorsal organ warming cells (DOWCs)(*25*, *26*). These neurons use distinct ionotropic receptors (IRs) to detect opposite signs of temperature change: DOCC cooling sensitivity requires *Ir21a*, whereas DOWC warming sensitivity requires *Ir68a*; both pathways require the IR co-receptors *Ir25a* and *Ir93a*(*25–30*). Larvae regulate navigation along thermal gradients primarily through turning rate: they suppress turning when a temperature change moves them toward their preferred temperature and increase turning when it moves them away(*26*). How larvae assign appropriate behavioral valence to warming and cooling around a preferred setpoint has been poorly understood.

The physiological properties of DOCCs and DOWCs create a core puzzle relevant to thermoregulation in many systems. While these peripheral pathways report the direction of temperature change, they operate similarly throughout the innocuous range, both above and below the preferred temperature. Importantly, they do not by themselves encode absolute temperature or specify the setpoint(*25*, *26*, *29*). Yet the behavioral valence of the temperature-change signals they transmit depends on absolute temperature: cooling is avoided below the preferred setpoint and warming is avoided above it. Thus, while DOCCs and DOWCs provide exquisite sensitivity to warming and cooling (e.g., DOCCs detect ≤0.005°C /sec change(*25*)), their activity does not explain how thermosensory signals determine the behavioral setpoint.

Orthologs of the Transient Receptor Potential ion channel TRPA1 have been implicated in thermosensing in both invertebrates and vertebrates(*13*, *31–33*). In adult *Drosophila*, *TrpA1* mediates warmth activation of thermosensors within the fly head and is required for normal thermal preference behavior(*32*), but how it exerts this effect at a circuit level is not well understood. TRPA1 is also expressed in the larval brain(*27*), but its role in larval thermoregulation at innocuous temperatures is unknown(*32*).

Here, we define a small set of *TrpA1*-dependent core thermosensors, the Lateral Posterior Brain Thermosensors (LPBTs), that link the activity of peripheral warming and cooling pathways to an absolute temperature-dependent turning decision. The LPBTs lie two synapses downstream of the dorsal organ thermosensors and express the highly thermosensitive TRPA1 isoform TRPA1(A). LPBTs provide a crucial site for thermosensory integration in the central brain, as LPBT activity is necessary for larvae to reverse their response to warming above the preferred temperature, and LPBT activation suffices to elicit turning. Despite expressing the warmth-activated cation channel TRPA1(A), LPBTs are not standalone warmth sensors. Rather, they use TRPA1(A) in a gain control mechanism, amplifying incoming peripheral warming signals only when the internal absolute temperature exceeds ∼25°C. This work provides a new model for behavioral thermoregulation, in which core thermosensors tune the behavioral valence of peripheral thermosensory input.

## Results

### TRPA1 tunes warming behavior around the thermotactic setpoint

Previous work showed that, in *Drosophila* larvae, TRPA1 acts in peripheral nociceptors to drive avoidance of noxious heat (≥40°C)(*21*, *22*, *27*, *34*, *35*). Here, we study innocuous temperatures between 20°C and 30°C, where larvae navigate up or down gradients toward a preferred intermediate body temperature setpoint. In this temperature range, both positive and negative thermotaxis (movement up and down the gradient, respectively) require dorsal organ cooling cells (DOCCs) and dorsal organ warming cells (DOWCs)(*25*, *26*, *29*, *30*). DOCCs and DOWCs are activated by cooling and warming, respectively, across this range, both above and below the preferred setpoint. As thermotaxis toward the setpoint requires that behavioral output downstream of DOCCs and DOWCs is tuned by an additional mechanism sensitive to absolute temperature, we asked whether TRPA1 might contribute such information.

To assess how absolute temperature modulates behavioral responses to temperature change, we exposed freely crawling larvae to sinusoidal temporal oscillations of temperature around a range of median temperatures (*T_mid_*) while simultaneously tracking navigational behavior (Fig. 1A)(*25*, *26*). As previously described, navigational behavior was segmented into an alternating sequence of turns and runs. With fixed oscillation amplitude, turning depended strongly on *T_mid_*. Turning increased during warming at warmer median temperatures and during cooling at cooler median temperatures (Fig. 1B,C), responses which drive the larva toward its preferred body temperature. This imbalance between warming- and cooling-evoked turning increased with distance from the preferred setpoint (*T_pref_*) (Fig. 1B, solid lines).

**Fig. 1.**
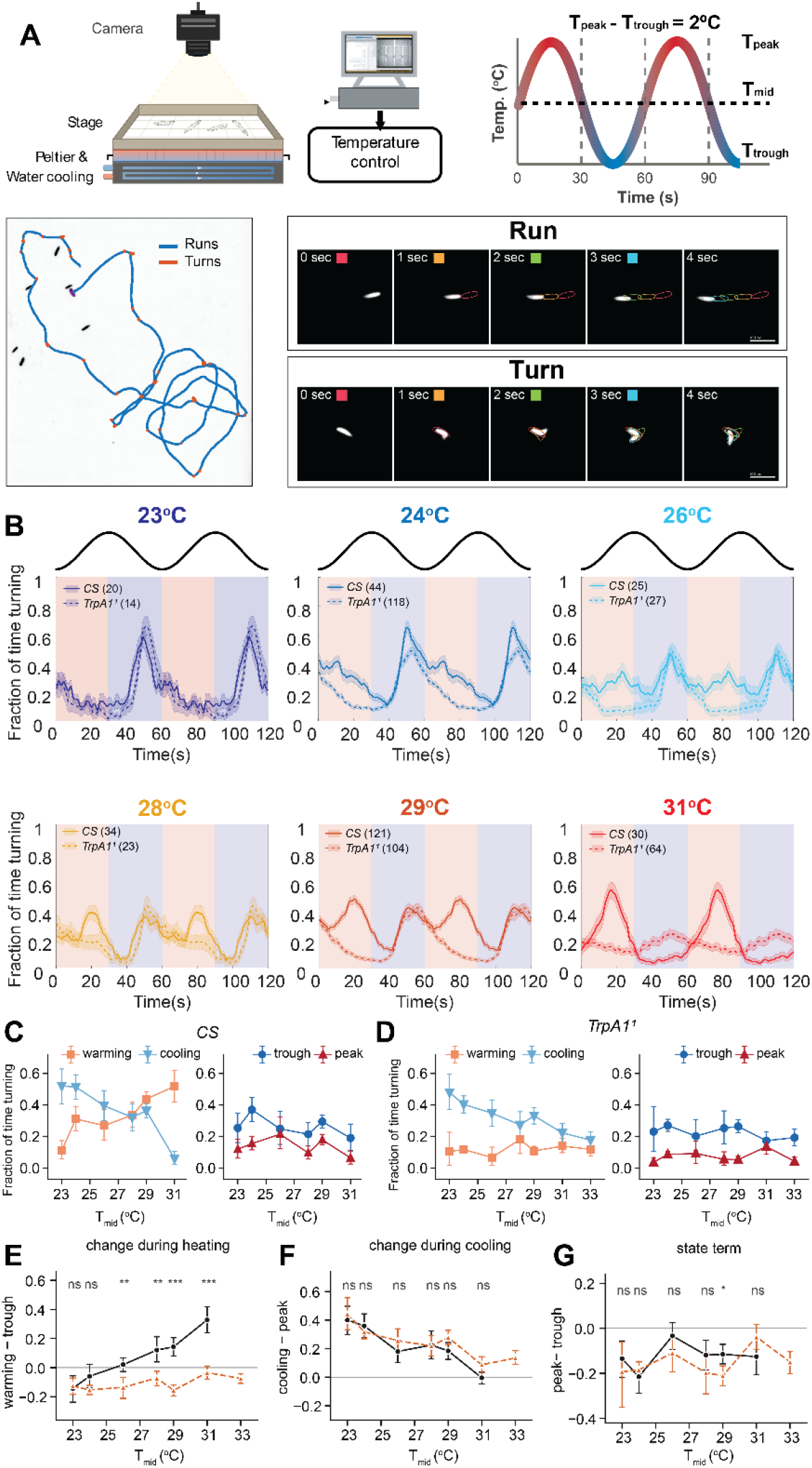
Turning responses to temperature oscillation depend on median temperature, and the warming response requires *TrpA1*. **(A)** Behavioral assay. Freely crawling larvae on an agar-covered stage were imaged from above while stage temperature was controlled by a Peltier element with water cooling under closed-loop feedback. Temperature was oscillated sinusoidally with a period of 60 s and an amplitude of ±1 °C about a median temperature (*T_mid_*), giving a peak (*T*_peak_) and trough (*T*_trough_) 2 °C apart. Bottom left, example trajectory of a single larva with behavior segmented into runs (blue) and turns (orange). Bottom right, representative image sequences of a run and a turn at 1 s intervals; colored outlines show the fitted body contour over successive frames. **(B)** Fraction of time spent turning averaged over one oscillation cycle repeated twice for Canton-S (*CS*, solid) and *TrpA1^1^* (dashed) at six median temperatures. Shading indicates the warming (pink) and cooling (blue) half-cycles; the stimulus waveform is drawn above each panel. Lines and bands show mean *±* SEM; *n* larvae per genotype is given in parentheses. In controls, turning is concentrated in the cooling phase at low *T_mid_* and shifts to the warming phase as *T_mid_* rises. In *TrpA1^1^*, the cooling-phase response is retained while the warming-phase response fails to emerge. **(C, D)** Mean fraction of time turning in each quarter-cycle as a function of *T_mid_*, for Canton-S **(C)** and *TrpA1^1^* **(D)**. Left, the two ramp phases (warming, cooling); right, the two static phases (trough, peak). Points show the mean and error bars the 95% CI. **(E–G)** Within-animal contrasts derived from the phase means. **(E)** Change during warming, defined as turning during warming minus turning at the trough. **(F)** Change during cooling, defined as turning during cooling minus turning at the peak. **(G)** State term, defined as turning at the peak minus turning at the trough. Canton-S in black, *TrpA1^1^* in orange. The warming contrast increases with *T_mid_* in controls and crosses zero at *T_mid_* = 27.4 °C (95% CI 27.0 °C to 27.9 °C), but remains flat and negative in mutants across the entire range tested **(E)**, whereas the cooling contrast **(F)** and the state term **(G)** are similar between genotypes. Points show the mean and error bars bootstrap 95% confidence intervals (20,000 resamples). Markers denote comparison between genotypes at each *T_mid_* (Mann–Whitney *U*, two-sided, Holm–Bonferroni corrected): *** *p <* 0.001, ** *p <* 0.01, * *p <* 0.05, *ns* not significant. The replication unit is the individual larva.

In wild-type (Canton-S) animals, warming- and cooling-evoked turning balanced near *T_mid_* = 27.4°C (95% CI 27.0 to 27.9, from linear fits of Fig.1C), considered here the operational *T_pref_*. *TrpA1* mutants behaved differently (Fig. 1B, dashed lines). While cooling-evoked turning in *TrpA1* mutants resembled that in wild-type animals, warming-evoked turning did not increase with *T_mid_*, with the difference becoming significant around 25°C (Fig. 1D,E). Turning during cooling declined with *T_mid_* in both genotypes and did not differ significantly between them at any *T_mid_* (Fig. 1F). The difference between turning at the peak and trough of each sinusoidal stimulus was likewise similar between genotypes (Fig. 1G). A second, independently derived *TrpA1* allele, *TrpA1^ins^*, had a similar phenotype, while its *w^1118^* background control behaved like Canton-S (Fig. S1). Thus, *TrpA1* loss selectively abolishes the increase in warming-evoked turning observed at warmer absolute temperatures, without broadly impairing thermosensitivity or turning.

### The warmth-activated thermosensor TRPA1(A) is necessary and sufficient for warming-evoked turning above setpoint

As *TrpA1* encodes multiple isoforms with different sensory properties, understanding *TrpA1*’s role in warming-evoked turning requires first establishing which of *TrpA1*’s functionally distinct isoform(s) are involved(*22*, *36*, *37*). Among the primary isoforms, TRPA1(A) and TRPA1(B) are expressed primarily in the larval central nervous system and exhibit warmth-sensitivity, while TRPA1(C) and TRPA1(D) are expressed primarily in class IV multidendritic neurons and mediate responses to damaging stimuli including noxious chemicals and noxious heat (≥40 °C)(*22*, *36*, *37*) (Fig. 2A).

**Fig. 2.**
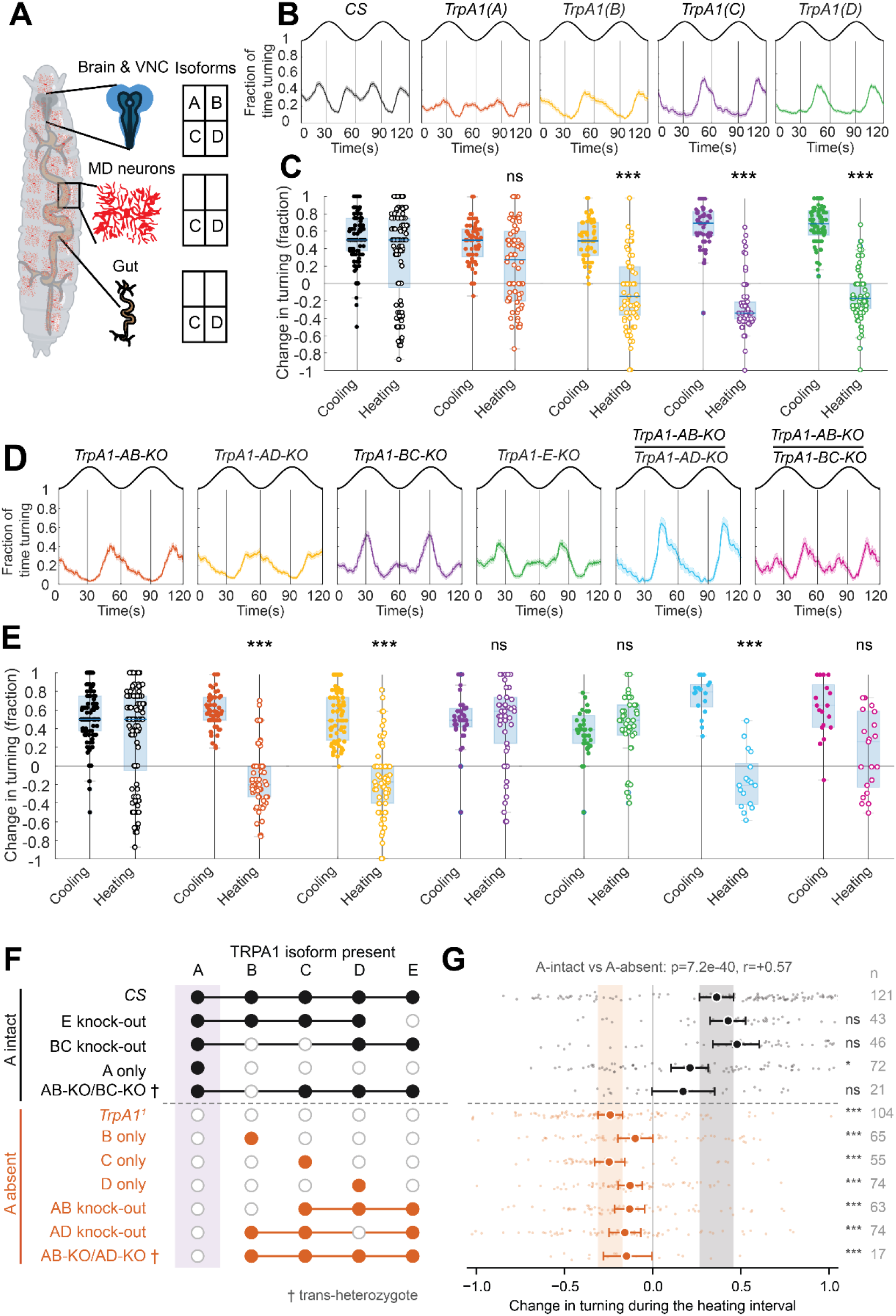
The A isoform of TRPA1 is necessary and sufficient for the warming-evoked turning response. **(A)** Reported expression of the four principal TRPA1 isoforms in the larva. Isoforms A, B, C and D are all expressed in the brain and ventral nerve cord, whereas only C and D are found in the class III and class IV multidendritic (MD) neurons of the body wall and in the gut. **(B)** Fraction of time spent turning across averaged one temperature oscillation cycle with *T_mid_* = 29°C repeated twice for Canton-S (*CS*) and for larvae expressing a single TRPA1 isoform. Vertical lines mark the boundaries between warming and cooling half-cycles. Lines and bands show mean *±* SEM. **(C)** Change in turning during the cooling (filled symbols) and warming (open symbols) intervals for the genotypes in **(B)**. Circles are individual larvae; boxes show the median and interquartile range. Only larvae expressing isoform A increase turning during warming; those expressing B, C or D alone show a decrease, as does the *TrpA1*^1^ null. The cooling response is unaffected in every genotype. **(D)** Turning across averaged one temperature oscillation cycle with *T_mid_* = 29°C repeated twice for larvae in which pairs of isoforms have been knocked out, and for the two trans-heterozygous combinations. The stimulus waveform is drawn above each panel in **(B)** and **(D)**. **(E)** Change in turning during cooling and warming for the genotypes in **(D)**. Removing isoform A together with either B or D abolishes the warming response, whereas removing B and C together, or removing E alone, leaves it intact. **(F)** Functional isoform composition of every genotype tested. Filled circles indicate an isoform that is present; open circles indicate one that is absent. Genotypes retaining A (black, above the dashed line) are separated from those lacking it (orange, below), with *CS* and *TrpA1*^1^ heading their respective groups as references. *†* denotes trans-heterozygotes, in which an isoform is scored as present if it is intact on at least one chromosome. **(G)** Change in turning during the warming interval at *T*_mid_ = 29°C for the genotypes in **(F)**. Small dots are individual larvae, jittered; circles show the mean with bootstrap 95% confidence intervals (20,000 resamples). Shaded vertical bands mark the confidence intervals of *CS* (grey) and *TrpA1*^1^ (orange). Every genotype retaining isoform A falls within or adjacent to the *CS* band and every genotype lacking it falls within or adjacent to the *TrpA1*^1^ band, irrespective of which other isoforms are present. Pooled across genotypes, A intact (+0.340, *n* =303 larvae) differs from A absent (*−*0.170, *n* =452; *p* =7.2*×*10*^−^*^40^, Mann–Whitney *U*, rank-biserial *r* =+0.57). The two trans-heterozygotes constitute a complementation test: *AB*-KO/*BC*-KO retains one functional copy of A and is indistinguishable from *CS* (*p* =0.056), whereas *AB*-KO/*AD*-KO lacks A on both chromosomes and differs from *CS* (*p* =3.5*×*10*^−^*^4^), despite sharing the *AB*-KO chromosome with the former. Throughout, asterisks denote comparison of each genotype with *CS* (Mann– Whitney *U*, two-sided): *** *p <* 0.001, ** *p <* 0.01, * *p <* 0.05, ns not significant. *n* larvae per genotype is given at the right of **(G)**; the replication unit is the individual larva.

To probe which *TrpA1* isoforms contribute, we assayed warming-evoked turning at *T_mid_* = 29°C, an innocuous temperature at which *TrpA1* mutants differ significantly from wild type. To identify the isoform(s) sufficient to support the behavior, we examined animals in which the *TrpA1* locus has been engineered to express only single isoforms(*37*). Consistent with the TrpA1(A) isoform driving this behavior, TRPA1(A)-expressing larvae exhibited warming-evoked turning at *T_mid_* = 29°C, while larvae expressing other isoforms showed low turning or decreased turning during the warming phase (Fig. 2B,C). To test which isoforms are necessary, we examined mutants in which the *TrpA1* locus has been engineered to eliminate the production of pairs of isoforms (Fig. 2D,E)(*37*). Further consistent with a critical role for the A isoform, animals lacking the A isoform, *TrpA1-AB-KO* and *TrpA1-AD-KO* mutants, phenocopied *TrpA1* mutants, while *TrpA1-BC-KO* and *TrpA1-E-KO* mutants did not (Fig. 2D,E). In addition, *TrpA1-AB-KO/TrpA1-AD-KO* animals, in which only the A isoform is homozygous mutant, also phenocopies *TrpA1* mutants, further confirming the TRPA1(A) is both necessary and sufficient to support warming-evoked turning at temperatures above the setpoint (Fig. 2D,E,F,G). TRPA1(A) functions as a warmth-activated cation channel, commonly eliciting neuronal activation at ≥25°C(*32*, *36*). This further suggests that neurons expressing TRPA1(A) mediate warming-evoked turning above setpoint.

### LPBTs drive TRPA1-dependent warming-evoked turning

We next sought to identify which TRPA1(A)-expressing neurons contribute to the homeostatic setpoint. TRPA1 isoforms are expressed in distinct but partially overlapping subsets of neurons (Fig. 2A). Isoforms C and D are expressed outside the central nervous system, including in multidendritic neurons and the midgut, whereas isoforms A and B are restricted to the brain, Ventral Nerve Cord (VNC) and corpus cardiacum. The A isoform is expressed in approximately 40 to 45 neurons in the brain and 12 neurons in the VNC.

RNAi knockdown of *TrpA1* in either all *TrpA1*-expressing cells or A-isoform-expressing neurons reproduced the *TrpA1* mutant’s warming-evoked turning phenotype (Fig. 3A,B). In contrast, knockdown of *TrpA1* in the corpus cardiacum, where TRPA1(A) is also expressed, with *Akh-GAL4* did not. Knockdown with the C-isoform driver *TrpA1(C)-GAL4* also reproduced the mutant phenotype. Because *TrpA1(C)-GAL4* labels a small number of neurons in the brain and VNC, the overlap between *TrpA1(C)-GAL4*-labelled and TRPA1(A)-expressing neurons narrowed the candidate population to three neurons per hemisphere, which we named Lateral Posterior Brain Thermosensors (LPBTs) (Fig. 3C,D).

**Fig. 3.**
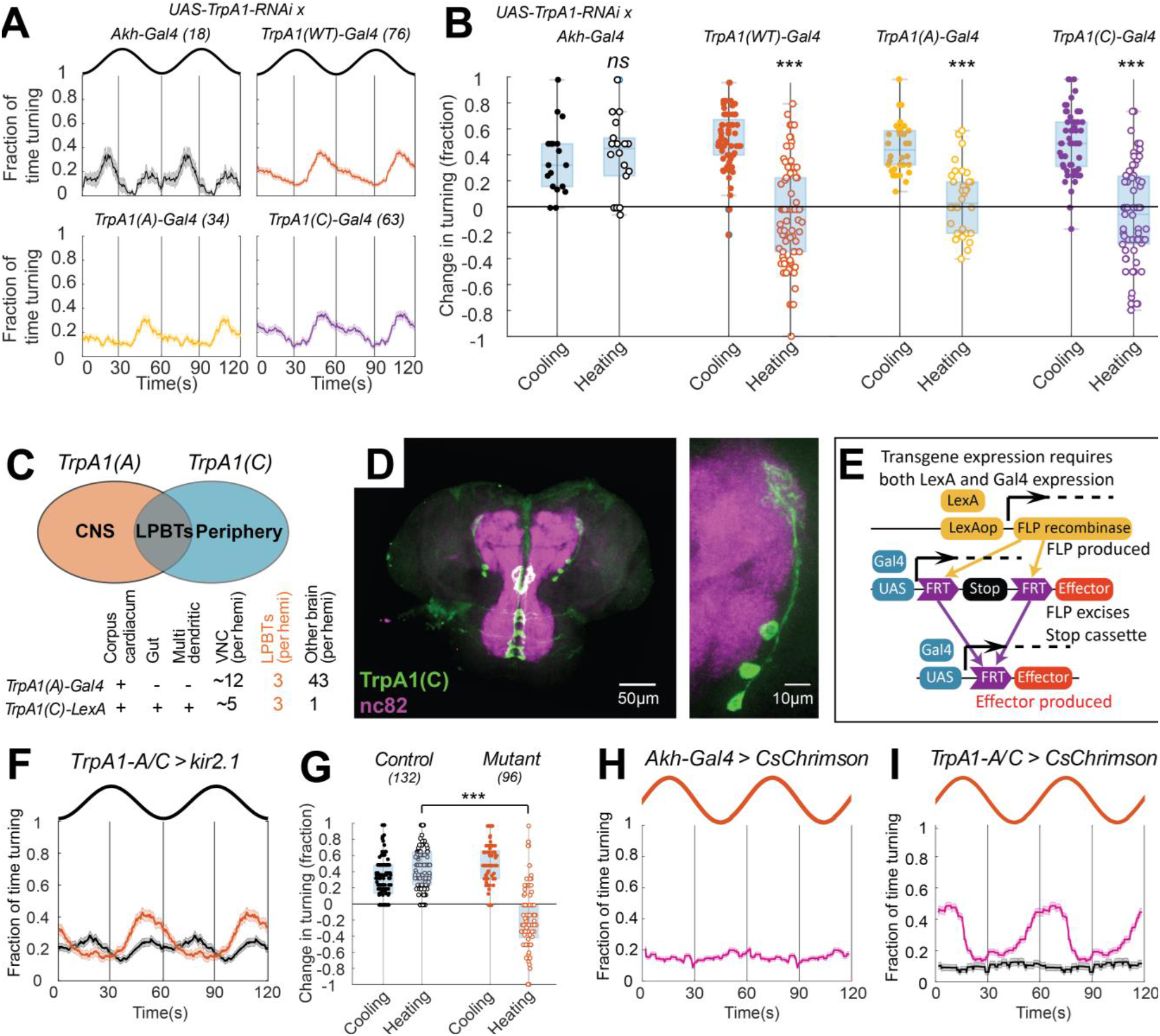
LPBTs in the central brain control the behavioral response to warming. (**A** and **B**) *TrpA1* knocked down by RNAi under four drivers, assayed at *T*_mid_ = 29 °C: fraction of time turning across two oscillation cycles (**A**) and change in turning during cooling and heating (**B**). The stimulus waveform is shown above each panel in (**A**), and *n* larvae are given in the brackets. Knockdown with *TrpA1(WT)-GAL4*, *TrpA1(A)-GAL4* or *TrpA1(C)-GAL4* abolished the increase in turning during heating (all *p* ≤ 4 × 10⁻⁴ against Canton-S), whereas knockdown restricted to the corpus cardiacum with *Akh-GAL4* did not (*p* = 0.54). Cooling responses were unaffected in all four genotypes. (**C**) Overlap between the *TrpA1(A)-GAL4* and *TrpA1(C)-LexA* expression patterns. *TrpA1(A)* is expressed predominantly in the central nervous system and *TrpA1(C)* predominantly in the periphery; the two intersect in three neurons per hemisphere, the LPBTs. The table gives expression per tissue and the number of labeled cells per hemisphere. (**D**) Maximum intensity projections of a larval central nervous system in which *TrpA1(C)-GAL4* drives GFP (green), counterstained with nc82 (magenta). Left, brain and a piece of ventral nerve cord; right, one hemisphere at higher magnification showing the LPBT cell bodies and their projection. Scale bars, 50 µm and 10 µm. (**E**) Intersectional strategy used to restrict transgene expression to LPBTs. *TrpA1(C)-LexA* drives FLP recombinase, which excises an FRT-flanked transcriptional stop cassette from a *TrpA1(A)-GAL4*–driven effector, so that the effector is produced only in neurons positive for both drivers. (**F** and **G**) Silencing LPBTs with Kir2.1 (orange) abolished the heating response while leaving the cooling response intact; Control (black), the same cross without *UAS-Kir2.1*. (**H** and **I**) Optogenetic activation. The sinusoidal trace shows red light intensity in arbitrary units, delivered with the same period as the thermal stimulus. CsChrimson expressed in the corpus cardiacum did not modulate turning (**H**), whereas CsChrimson in LPBTs drove turning in phase with the light (magenta), absent in animals reared without ATR (black) (**I**). Lines and bands show mean ± SEM; circles in (**B**) and (**G**) are individual larvae, and boxes show the median and interquartile range with whiskers extending to the most extreme observation within 1.5 times the interquartile range. Asterisks denote comparison with Canton-S (**B**) or between the two genotypes (**G**) by two-sided Mann– Whitney *U* test with Holm–Bonferroni correction: \*\*\**p* < 0.001, ns, not significant.

We used an intersectional FLP-recombinase strategy to exploit the narrow overlap between *TrpA1(A)-GAL4* and *TrpA1(C)-LexA*, restricting effector expression almost exclusively to LPBTs(*38*, *39*). In this strategy, *TrpA1(C)-LexA* drove *LexAop-FLP*, while *TrpA1(A)-GAL4* drove *UAS-FRT-stop-FRT-[effector]* (Fig. 3E). The transcriptional stop cassette flanked by FRT sites is excised by FLP only in cells where the *LexA* driver is active, thereby restricting effector expression to neurons positive for both the *GAL4* and *LexA* drivers. Silencing LPBTs with this approach reproduced the *TrpA1* mutant phenotype: larvae failed to increase turning during warming while their cooling response remained intact (Fig. 3F,G). We also tested whether LPBT activation is sufficient to drive turning, the behavioral output that increases during warming above the setpoint. Optogenetic activation of LPBTs with CsChrimson drove rapid turning when the sinusoidal temperature oscillation was replayed as red-light intensity (Fig. 3H,I). LPBTs are therefore necessary for the warming-evoked turning response and sufficient to elicit turning when activated.

### LPBT temperature responses have both TRPA1-dependent and TRPA1-independent components

To characterize LPBT responses to temperature, we used the genetically encoded calcium indicator GCaMP7b. Larvae were first held at 18°C, then exposed to a 1°C warming ramp at 0.1°C/sec, a 20s hold, and a cooling ramp at the same rate (Fig. 4A). LPBT neurites exhibited sharp increases at the onset of warming and maintained an elevated plateau throughout the hold, followed by rapid decreases during cooling (Fig. 4B,C). In contrast, LPBT cell bodies responded more gradually during warming, with fluorescence increasing across successive temperature steps. Thus, LPBT responses are not simply monotonic increases to increased temperature as one might predict from a simple warmth sensor. Instead, the differences between cell body and neurite responses suggested LPBTs might integrate temperature signals in a more complex fashion.

**Fig. 4.**
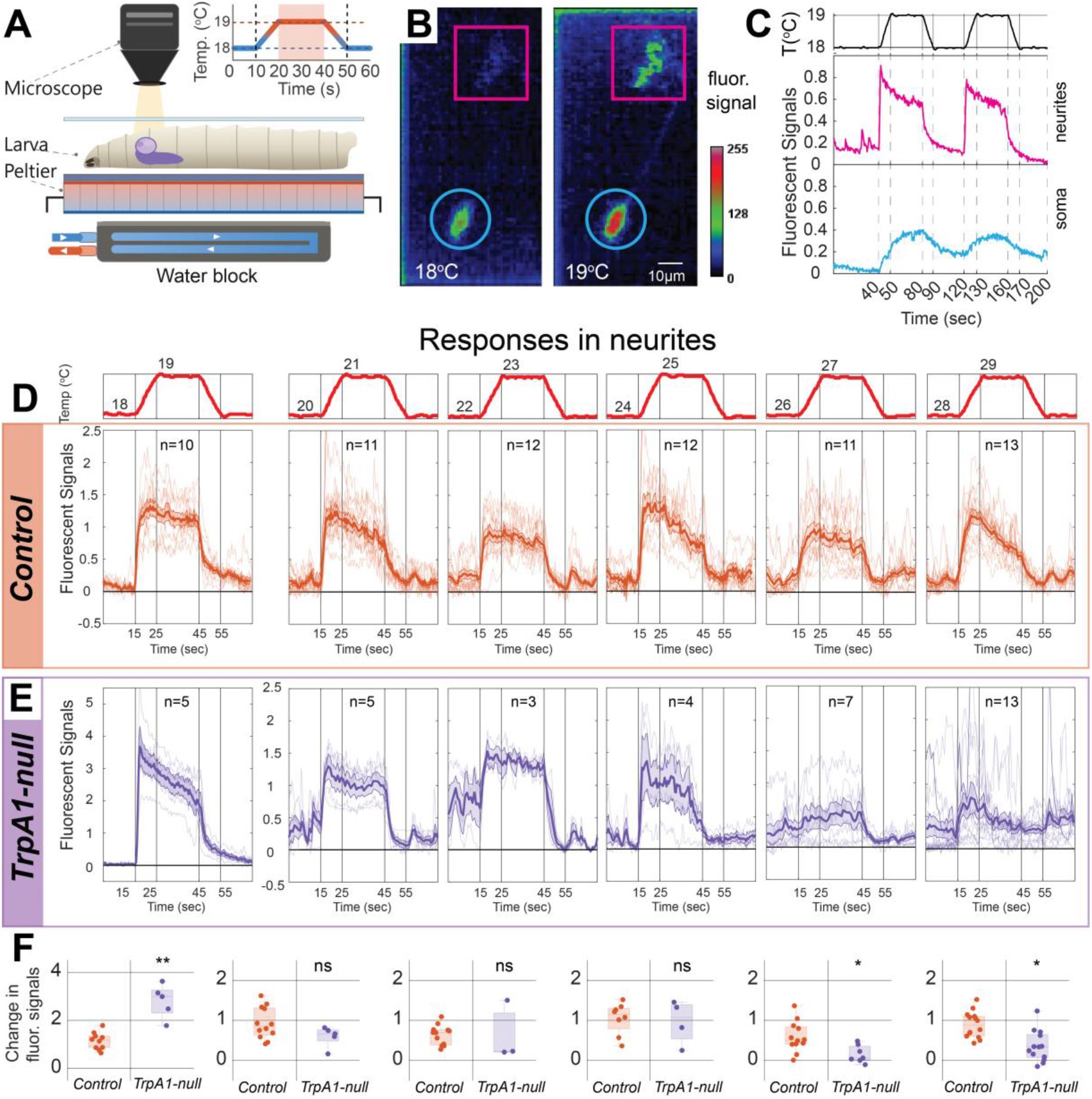
LPBTs respond to warming across a wide temperature range, and the contribution of TRPA1 is itself temperature dependent. **(A)** Imaging preparation. An semi-immobilized larva was mounted on a temperature stage with Peltier element and water cooling and imaged from above. Inset, temperature protocol: a 1 °C ramp delivered at 0.1 °C/sec, held for 20 s (shaded) and returned at the same rate. **(B)** Pseudocolored fluorescence of an LPBT at the base temperature (18 °C) and at the top of the ramp (19 °C), showing the cell body (cyan circles) and its neurites (magenta squares). Scale bar, 10 µm. **(C)** Stage temperature (top) and fluorescence in the neurites (magenta) and cell body (cyan) of a single LPBT during two successive 1 °C ramps. The neurite signal rises sharply at the onset of each ramp and decays during the plateau, whereas the somatic signal builds gradually and does not return to baseline between steps. **(D, E)** Fluorescence responses in neurites to 1 °C ramps delivered from six base temperatures (18 °C, 20 °C, 22 °C, 24 °C, 26 °C, and 28 °C; stage temperature shown in red above **D**). **(D)** Control larvae expressing GCaMP7b under *TrpA1(A)-GAL4* control. **(E)** *TrpA1-null* larvae expressing GCaMP7b under *TrpA1(C)-GAL4* control. The *TrpA1(A)-GAL4* driver was used for control to produce a similar expression levels of two alleles in a wild type animals. *TrpA1(C)-G*AL*4* driver was crossed with the *UAS-GCaMP7b*;*TrpA1*^1^ to completely eliminate TRPA1(A) gene. Responses in the animals expressing GCaMP7b with *TrpA1(C)-GAL4* (Fig.S3A) therefore with only one allele of *TrpA1(A)* are already reduced at the temperatures T_mid_ ≥ 24 °C Thin lines are individual animals; thick lines and bands show mean *±* SEM. **(F)** Change in fluorescence during the ramp for the genotypes in **(D)** and **(E)** at each base temperature. *n* indicates the number of animals per condition for D–F. Circles are individual animals; boxes show the median and interquartile range. Responses differed between genotypes at 18 °C, 26 °C, and 28 °C, but not at 20 °C, 22 °C, or 24 °C. Asterisks denote comparison between genotypes at each base temperature: *** *p <* 0.001, ** *p <* 0.01, * *p <* 0.05, *ns* not significant.

To establish how LPBTs respond to warming at different absolute temperatures, we subjected larvae to 1°C warming ramps delivered from different base temperatures. Since the calcium responses in neurites were stronger and somatic responses followed, we focused on analyzing the former. LPBT neurites showed sustained warming responses from base temperatures of 18°C through 29°C (Fig. 4D, Fig. S2A). Across this range, response amplitudes were comparable and activity persisted until the onset of cooling. Below 18°C, LPBT neurites did not respond to warming or responded but did not sustain activity through the hold, returning to baseline when warming ended (Fig. S2B). As these low-temperature responses occur well below temperatures at which TRPA1(A) activity has been observed, they suggested that LPBT warming responses may not solely depend on TRPA1.

We assessed the contribution of TRPA1 to LPBT thermosensing using *TrpA1* mutants. In these animals, LPBTs responses to warming remained robust at low temperatures, but were reduced or absent above ∼25°C (Fig. 4E). Below 24°C, mutant responses closely resembled controls, except at 18°C, where responses were stronger. For ramps initiated at 24°C, responses were not sustained at the 25°C peak. Finally, warming initiated from a base of 26 or 28°C produced substantially smaller LPBT responses than in wild type (Fig. 4E). Therefore, TRPA1 is not required for LPBTs to detect warming per se. Instead, *TrpA1* loss alters the LPBT response to warming specifically at elevated temperature: below ∼25°C, warming responses remain largely intact, whereas above 25°C, warming evokes little or no sustained activity, in the same temperature range as the behavioral phenotype (Fig. 1E).

Because LPBTs likely influence behavior through their synaptic output, we next asked how temperature integration is represented at their presynaptic terminals. We expressed a synaptotagmin-GCaMP6s fusion (syt-GCaMP6s) under *TrpA1(C)-GAL4* control, restricting the indicator to presynaptic terminals, and identified the LPBT terminal field in the lateral horn (Fig. S2C,D). Warming evoked a sharp rise in terminal calcium at the onset of the ramp, reaching Δ*f/f* =1.2. Unlike the sustained plateau recorded from all LPBT processes, the presynaptic terminal signal adapted during the hold, declining to approximately 20% of peak by the end of the plateau. A second, smaller transient followed the end of the cooling ramp likely due to a small temperature bump after the ramp ends. Thus the LPBT temperature response reaches the output compartment as a predominantly phasic signal that emphasizes warming onset.

The *TrpA1* mutation removes the channel from every cell. To ask whether TRPA1 is required within LPBTs themselves, we knocked it down with *TrpA1-RNAi* driven by *TrpA1(C)-GAL4*, which labels LPBTs and one additional neuron in the brain. LPBT calcium responses in these animals resembled those of the null mutant (Fig. S2E): warming evoked robust responses at 18 and 22°C, a reduced and poorly sustained response at 24°C, and at 26 and 28°C an initial transient that decayed rather than persisting through the plateau. The requirement for TRPA1 therefore maps to the sparse *TrpA1(C)-GAL4* brain population containing LPBTs, not to the peripheral dorsal organ thermosensors (DOCCs and DOWCs).

### LPBTs are downstream of dorsal organ ionotropic receptor pathways

The persistence of some LPBT temperature responses in the absence of *TrpA1* raised the possibility that other thermosensory pathways, like those from the periphery, might also provide temperature information to LPBTs. We therefore examined the larval connectome to assess potential connectivity between the peripheral dorsal organ thermosensors and LPBTs.

To identify LPBTs in the connectome, we used their fluorescently labeled position and morphology (Fig. 3) to search for matching neurons. This search identified three candidate neurons with LPBT-like morphology, located two synapses downstream of the dorsal organ thermosensors (Fig. 5A).

**Fig. 5.**
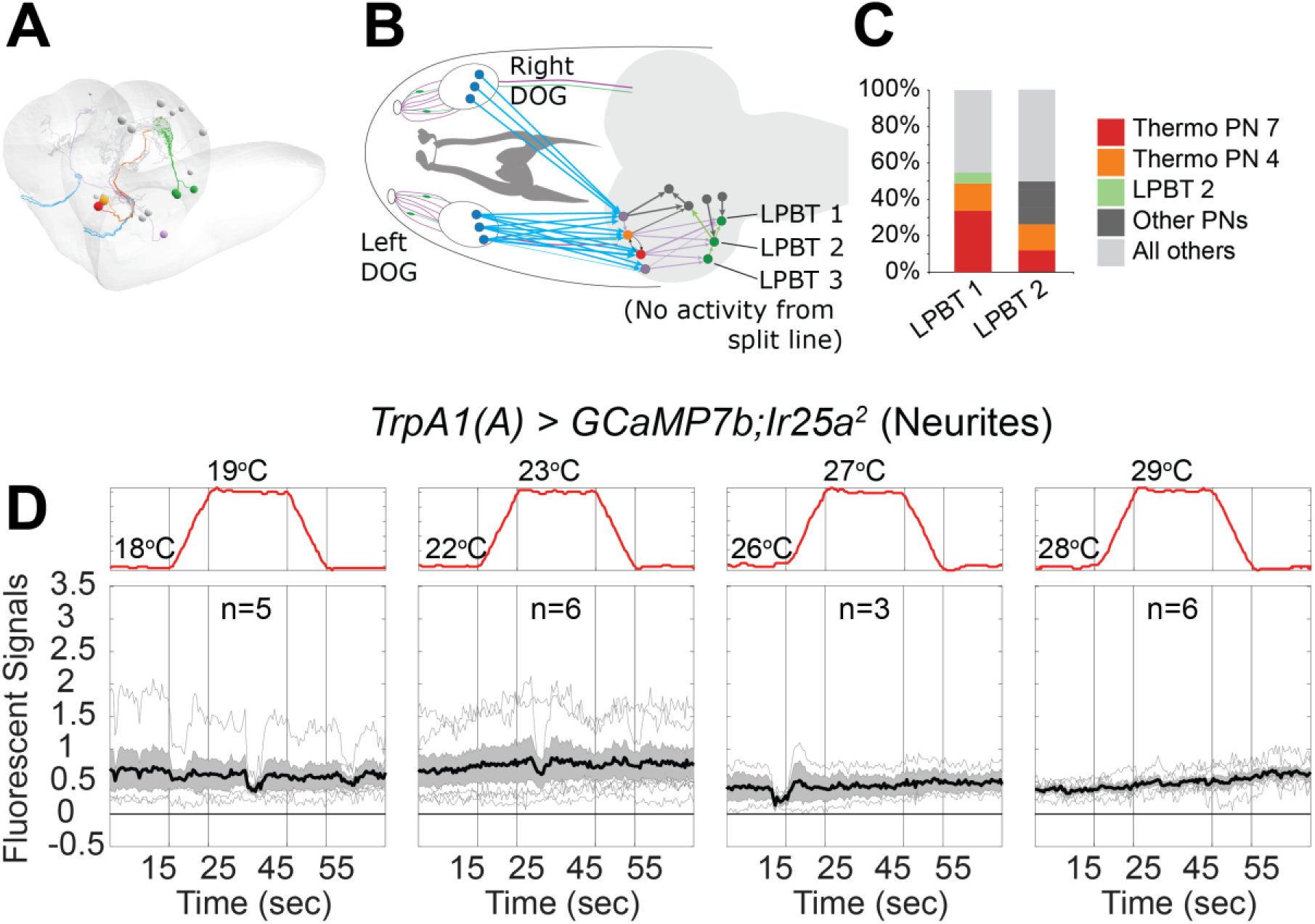
LPBT responses to temperature require peripheral thermosensory input. **(A)** Electron microscopy reconstruction of the three candidate LPBTs (green) and the thermosensory projection neurons that supply them, shown within the outline of the larval brain. **(B)** Connectivity from the left and right dorsal organ ganglia (DOG) to the LPBTs. Thermosensory receptor neurons (blue) project to thermosensory projection neurons tPN 7 (red), tPN 4 (orange), and others (grey), which in turn target LPBT 1, LPBT 2, and LPBT 3 (green). No temperature-evoked activity was recorded from the split-GAL4 line labeling LPBT 3. **(C)** Synaptic input to LPBT 1 and LPBT 2, expressed as a percentage of the total number of input synapses onto each cell. Thermosensory projection neurons 7 and 4 together account for 40% and 20% of input, respectively. **(D)** Fluorescence responses of LPBT neurites expressing GCaMP7b under *TrpA1(A)-GAL4* control in an *Ir25a*^2^ null background, during 1 °C ramps delivered from four base temperatures (stage temperature in red above each panel). Thin lines are individual animals; the thick line and band show mean *±* SEM. No temperature-evoked responses were detected at any base temperature.

We further confirmed this match using split-GAL4 lines that labeled individual LPBT candidate neurons and allowed us to measure single-cell temperature responses (Fig. S3). Of the three candidate neurons, two responded to temperature changes. These two neurons receive the majority of their input from projection neurons downstream of the dorsal organ thermosensors (Fig. 5B,C), suggesting that LPBTs receive peripheral temperature-change signals and transform them through a cell-intrinsic TRPA1-dependent mechanism.

To test whether dorsal organ thermosensors provide temperature-evoked drive to LPBTs, we disrupted their temperature responses by removing *Ir25a*, an ionotropic receptor co-receptor required for temperature responses in both dorsal organ cooling (DOCCs) and warming neurons (DOWCs). Measuring LPBT calcium responses in an *Ir25a*-null background (Fig. 5D), temperature-evoked responses were abolished in both somata and processes, even at 18°C, where LPBT responses are robust and do not require TRPA1 (Fig. S4A). Thus, LPBTs are functionally downstream of dorsal organ thermosensors, and are not entirely autonomous thermometers.

These findings demonstrate that LPBTs are not simple warmth receptors, but rather integrators of multiple thermosensory inputs: LPBTs combine input from peripheral thermosensors, reflecting environmental temperature change, with cell-intrinsic TRPA1-dependent thermosensory input, reflecting the absolute temperature inside the brain. In the LPBTs, we propose that TRPA1 acts to provide a temperature-dependent gain mechanism. This mechanism converts peripheral warming signals into warming-evoked turning as absolute temperature rises above the preferred range, thereby driving negative thermotaxis down thermal gradients and helping define the thermotactic setpoint.

## Discussion

There has been substantial progress in our understanding of the molecules and cells through which animals sense temperature. But how functionally diverse, spatially distributed thermosensory signals are integrated to facilitate body temperature homeostasis remains poorly understood. The small size and genetic accessibility of the *Drosophila* larval nervous system has allowed us to identify a circuit that combines input from peripheral pathways, reporting ambient temperature change, with that from core thermosensors, modulated by absolute temperature, to control body temperature homeostasis in the larva.

### Broadly tuned peripheral pathways interact with core thermosensors to set preferred temperature

The peripheral thermoreceptors of the dorsal organ ganglion (DOG) report the rate and direction of temperature change(*25*, *26*, *29*, *30*, *40*), providing larvae exquisite sensitivity to shallow (≤0.005°C/sec) thermal gradients. However, both DOCCs and DOWCs respond similarly across a broad range of innocuous temperatures spanning the preferred setpoint. Their physiological tuning therefore does not explain why cooling drives avoidance below the preferred temperature, why warming drives avoidance above it, or how an intermediate preferred body temperature setpoint is specified.

Here we begin to solve this puzzle by identifying a set of core brain thermosensors, the *TrpA1*-dependent LPBTs, that help process this peripheral input to drive the animal toward its preferred temperature. The LPBTs, which receive synaptic input from peripheral pathway neurons, are specifically required for larvae to reverse their response to warming as ambient temperature rises above the preferred setpoint. Below the setpoint, warming does not increase turning, allowing larvae to sustain runs up thermal gradients; above it, warming increases turning and biases movement down gradients. Silencing LPBTs abolishes this switch, disrupting the behavioral setpoint. Conversely, optogenetic activation of LPBTs is sufficient to elicit turning. These findings link thermosensory integration to the behavioral output that implements thermotaxis. Note that cooling-evoked turning below the setpoint persists in the absence of LPBT activity, raising the possibility that analogous pathways regulate these responses.

### TRPA1 tunes thermosensory gain without rendering LPBTs standalone warmth sensors

Beyond its normal functions, TRPA1(A) is a well-known tool for thermogenetic neuronal activation in *Drosophila*, as its ectopic expression can render neurons warmth-activated(*32*). In LPBTs, TRPA1(A) plays a distinct role. Rather than mediating LPBT warmth activation on its own, TRPA1(A) acts downstream of DOG peripheral thermosensory pathways to control the gain with which peripheral warming signals converging on the LPBTs are converted into behavior. Importantly, peripheral input suffices to elicit LPBT responses to warming even when TRPA1(A) is absent. However, TRPA1(A) expression is required to sustain LPBT warming responses once absolute temperature exceeds roughly 25°C, the same range in which warming-evoked turning becomes TRPA1(A)-dependent.

At a physiological level, LPBTs rely upon TRPA1(A) to amplify warming signals from the periphery at temperatures near and above the preferred setpoint. The compartmental organization of LPBT responses suggests how such a gain mechanism could work. LPBT neurites have sharp phasic responses at the onset of warming and are suppressed by cooling; terminals report the onset of warming but adapt during the plateau; and somatic signals build gradually and accumulate across successive temperature steps (Fig. 4). If a warming-evoked signal persists in LPBT neurites long enough to recruit the soma, a temperature-dependent conductance could make that recruitment conditional on absolute temperature. Thus, we would suggest that TRPA1(A) doesn’t carry sufficient current to initiate full-scale depolarization of the LPBTs on its own, but rather modulates the resting potential in an absolute-temperature-dependent manner to alter the spike threshold or excitability more generally. In this way, TRPA1 provides a temperature-dependent “gain control” that determine how strongly DOG-derived warming input is converted into LPBT output and warming-evoked turning above the preferred range.

### The gain control behavioral thermoregulation model

The TRPA1-dependent gain-control model suggested by our results argues against a view in which cool and warm receptors independently drive cold and heat avoidance. Rather than peripheral pathways acting alone, warming input is modulated by absolute temperature only two synapses downstream of the receptors. Interestingly, thermal preference in adult *Drosophila* also relies on a combination of IR-expressing peripheral thermosensors that primarily sense temperature change(*40*) and TRPA1-expressing internal thermosensors(*32*), raising the possibility that the strategy we describe in the larva may apply to adult flies. There are differences, as the heat-activated receptor Gr28b(D) required for heat avoidance is not expressed in larvae, suggesting a shared logic of integration and not necessarily its molecular implementation. Consistent with this, the switch in heat valence from avoidance to attraction in desert-dwelling *D. mojavensis* is associated not with the changes in peripheral receptors but with altered innervation of the lateral horn, the same region implicated in our work in larva(*41*).

It is interesting to speculate that TRPA1-dependent modulation of LPBT activity also provides a potential mechanism for changing the animal’s preferred temperature setpoint. The magnitude of current flowing through TRPA1 channels at a given temperature depends not only on the biophysics of individual channels, but on their level of expression. Alterations in TRPA1 channel abundance in LPBTs could shift the temperature at which the behavioral switch occurs without altering peripheral gradient detection, with increases in TRPA1 expression lowering the preferred temperature. Furthermore, the co-expression of a second isoform, TRPA1(C), in the LPBTs raises the possibility that altering the ratio of TRPA1 isoforms could also alter their activity. One can therefore imagine alterations in the levels and relative abundance of TRPA1 isoforms as a simple mechanism to adjust body temperature setpoints within *Drosophila melanogaster* in response to developmental or environmental stimuli, as well as in other *Drosophila* species with different thermal niches.

### Thermosensory integration across animals

Thermoregulatory circuits in other animals may use related molecular mechanisms and algorithms. In mammals, the preoptic area receives ascending thermal information from the periphery and coordinates diverse autonomic and behavioral thermoregulatory responses(*2*, *16*). A TRP-dependent mechanism similarly provides temperature-dependent amplification in the POA: TRPM2 acts presynaptically to make synaptic input onto preoptic warm-sensitive neurons temperature-dependent(*17*).

In summary, we have identified a larval circuit that explicitly links peripheral temperature-change signals, central absolute-temperature-dependent modulation, and motor control. LPBTs receive warming input from dorsal organ thermosensors, while TRPA1 acts within LPBTs to control how strongly that input is converted into LPBT output. In larval thermotaxis, modulation of a single behavioral control variable, turning rate, can bias navigation up or down thermal gradients. Because LPBT activation is sufficient to drive turning, this circuit directly links thermosensory integration to the single behavioral control variable that implements setpoint-directed navigation.

## Acknowledgments

We thank members of the Samuel and Garrity laboratories for helpful discussions.

## Author contributions

Conceptualization: SL, AS

Methodology: SL, AS, MZ, AC, PG

Investigation: SL

Visualization: SL

Funding acquisition: SL, AS, PG

Project administration: AS, PG

Supervision: AS, PG

Writing – original draft: SL, AS

Writing – review & editing: SL, AS, MZ, AC, PG

## Data and code availability

The data and code used for the manuscript will be made available through public repositories.

## Supplement Supplemental figures

**Fig. S1.**
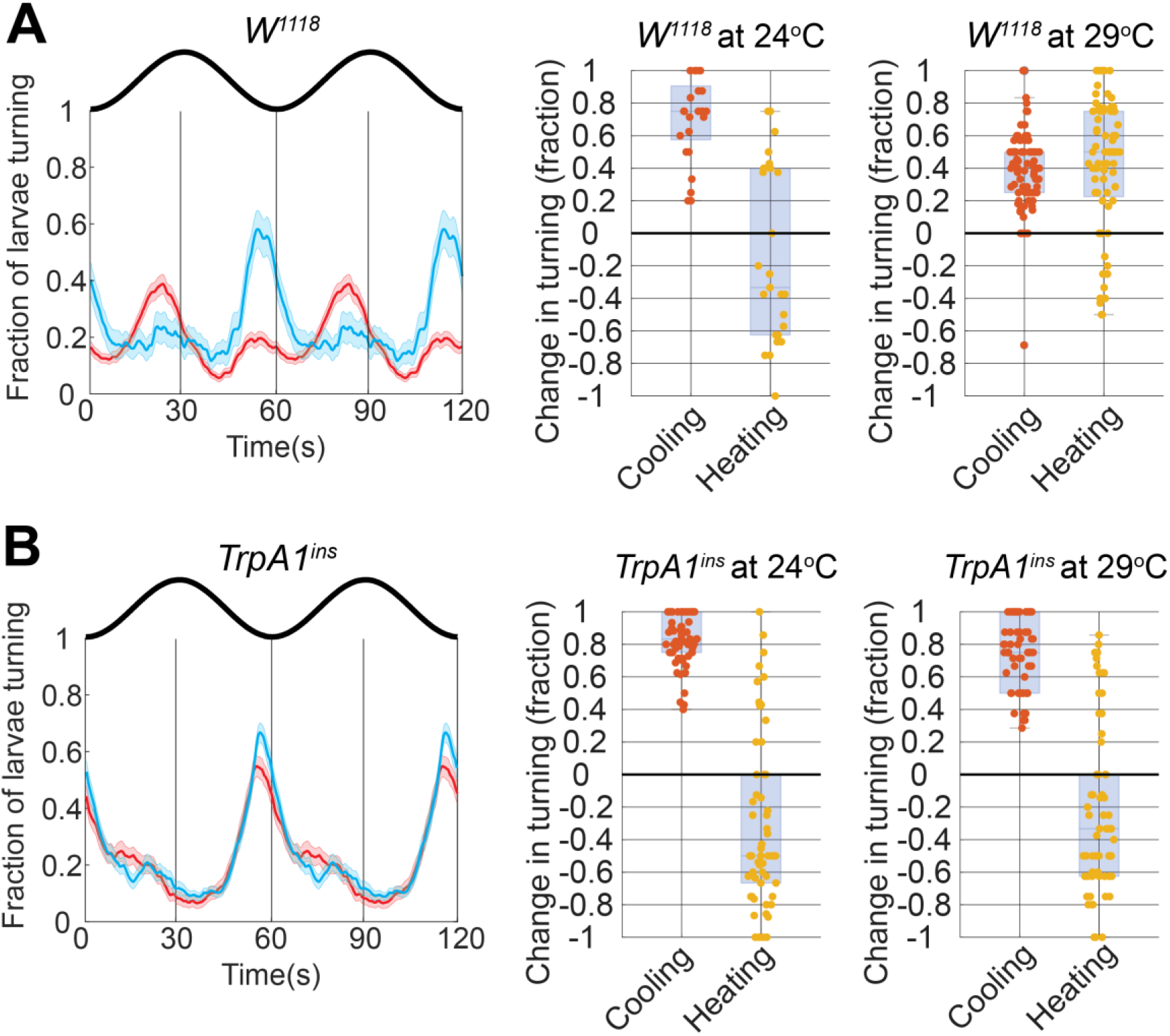
A second *TrpA1* allele reproduces the behavioral phenotype, while larvae mutant for the white gene do not. **(A)** *TrpA1^ins^* larvae. **(B)** *w^1118^* mutant. Left, fraction of larvae turning across two oscillation cycles at *T*_mid_ = 24°C (blue) and 29 °C (red), with the stimulus waveform above; lines show the mean and bands the SEM. Center and right, change in turning during the cooling and warming intervals at each median temperature. Circles are individual larvae; boxes show the median and interquartile range. *w^1118^* larvae reversed the sign of the warming response between 24 °C and 29 °C, as Canton-S does, whereas *TrpA1^ins^* larvae turned during cooling at both median temperatures and never increased turning during warming. The phenotype therefore follows the *TrpA1* lesion rather than the genetic background. (n=36 for *w^1118^* at *T*_mid_ = 24°C, n = 84 for *w^1118^* at *T*_mid_ = 29°C, n=59 for *TrpA1^ins^* at *T*_mid_ = 24°C, n=62 for *TrpA1^ins^* at *T*_mid_ = 29°C)

**Fig. S2.**
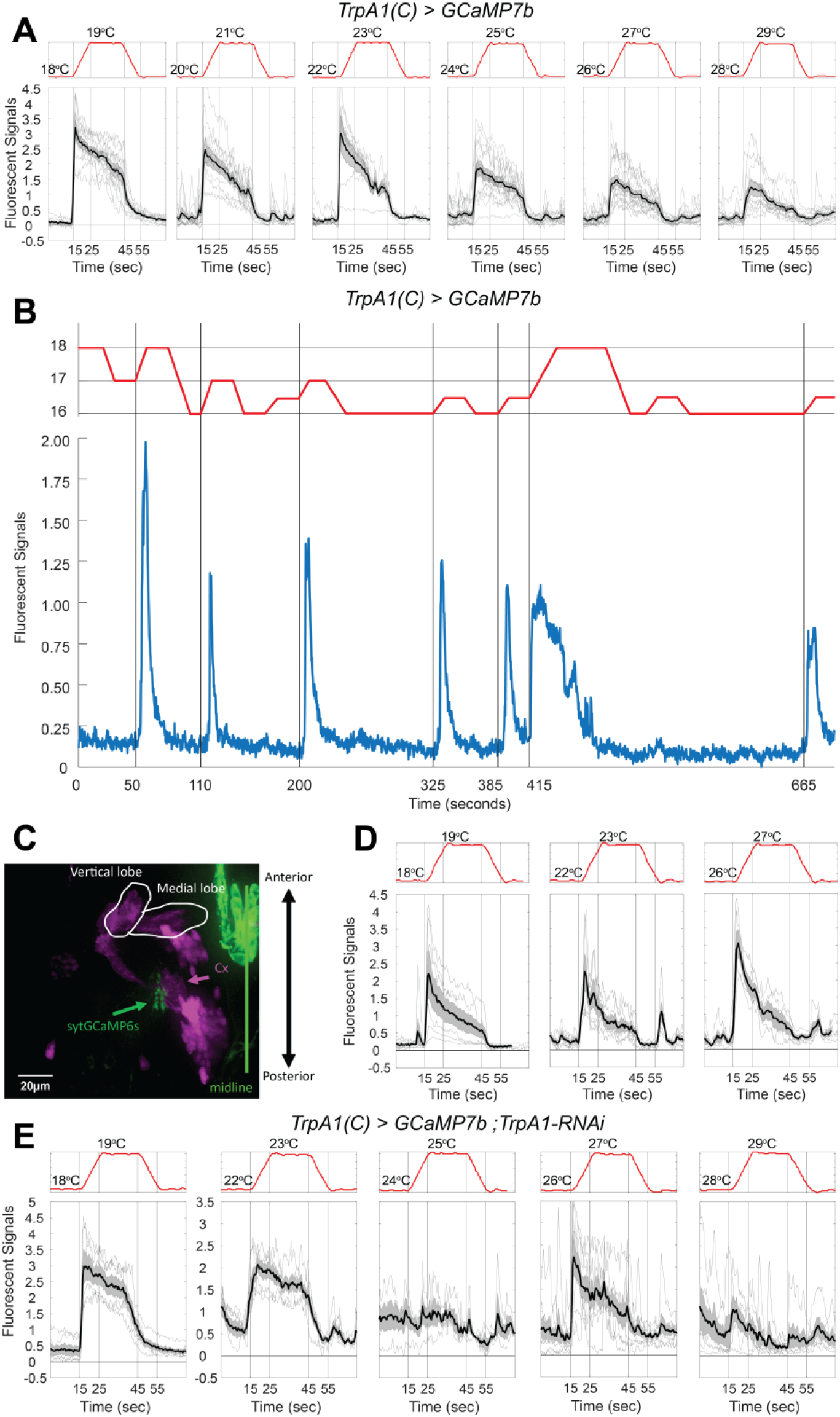
LPBT responses at various temperatures with *TrpA1(C)* driver, presynaptically expressed GCaMP and suppressed *TrpA1*. **(A)** Responses to 1 °C ramps from six base temperatures in larvae expressing *GCaMP7b* under *TrpA1(C)-GAL4* control. The responses show reduced amplitude and decrease during temperature plateau unlike the signals from *TrpA1(A)-GAL4* driver. Due to the nature of the transgenic mutations, the *TrpA1(C) > GCaMP7b* animals only have one functional *TrpA1(A)* allele, while *TrpA1(A) > GCaMP7b* animals, used in the Fig. 4D, have both. **(B)** Stage temperature (red) and fluorescence (blue) in a single LPBT expressing GCaMP7b under *TrpA1(C)-GAL4* control, during a sequence of 1 °C and 2 °C steps between 16 °C and 18 °C. Each increase in temperature evoked a transient response, and no response followed a decrease. **(C)** Expression of synaptotagmin-GCaMP6s (green) driven by *TrpA1(C)-GAL4* in the lateral horn, with the mushroom body labeled in magenta; the mushroom body vertical and medial lobes and the calyx (Cx) are indicated as landmarks. Scale bar, 10 µm. **(D)** Fluorescence responses in the LPBT terminal field expressing synaptotagmin-GCaMP6s to 1 °C ramps delivered from three base temperatures. **(E)** Responses to 1 °C ramps from five base temperatures in larvae expressing *TrpA1*-*RNAi* together with the calcium indicator under *TrpA1(C)-GAL4* control. Responses were retained at 18 °C and 22 °C but were reduced or absent from 24 °C upward, reproducing with cell-specific knockdown the temperature dependence seen in the *TrpA1*^1^ null (Fig. 4). In **(A, D, E)**, thin lines are individual animals and the thick line and shaded band show the mean and SEM; stage temperature is shown in red above each panel.

**Fig. S3.**
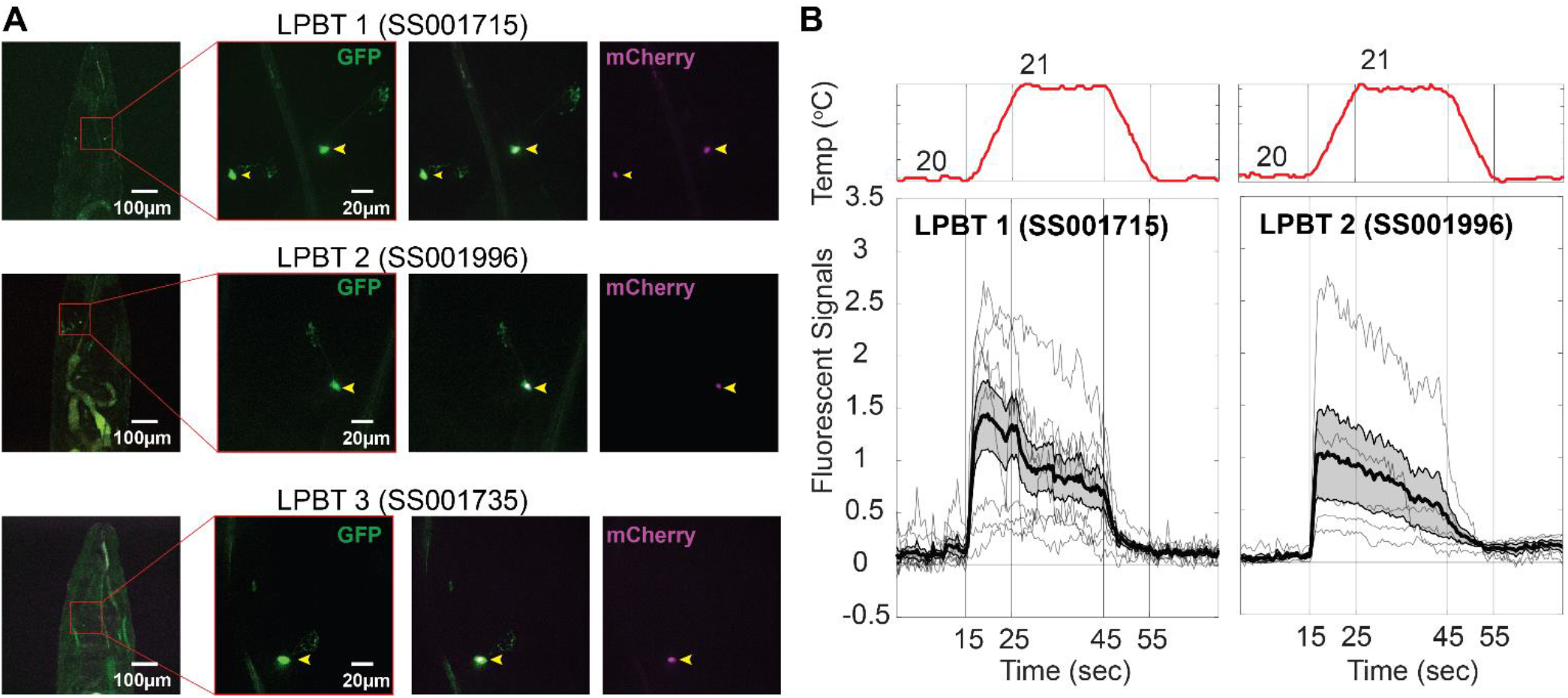
Split-GAL4 lines label LPBT neurons and reveal single-cell temperature responses. (**A**) Three split-GAL4 combinations, selected on the basis of the connectome reconstruction, each label one or more of the three LPBT candidates: SS001715 (R14E06-AD;R24E06-DBD, LPBT 1), SS001996 (R24E06-AD;VT045565-DBD, LPBT 2) andSS001735 (R24E06-AD;R14E06-DBD, LPBT 3). For each line, left, overview of a whole-mount second-instar larva with the boxed region shown at higher magnification to the right; then membrane bound GFP (green), the merged image, and nuclear mCherry (magenta). Yellow arrowheads mark the labeled cell bodies. Scale bars, 100 µm (overviews) and 20 µm (insets). (**B**) Fluorescence responses of LPBT 1 and LPBT 2 to a 1 °C warming ramp from 20 °C (stage temperature in red above each panel). Thin lines are individual animals; the thick line and shaded band show the mean and SEM. Both neurons responded with a sustained increase during warming that decayed during cooling. No temperature-evoked responses were detected in LPBT 3. n = 5, 8 and 5 animals respectively.

**Fig. S4.**
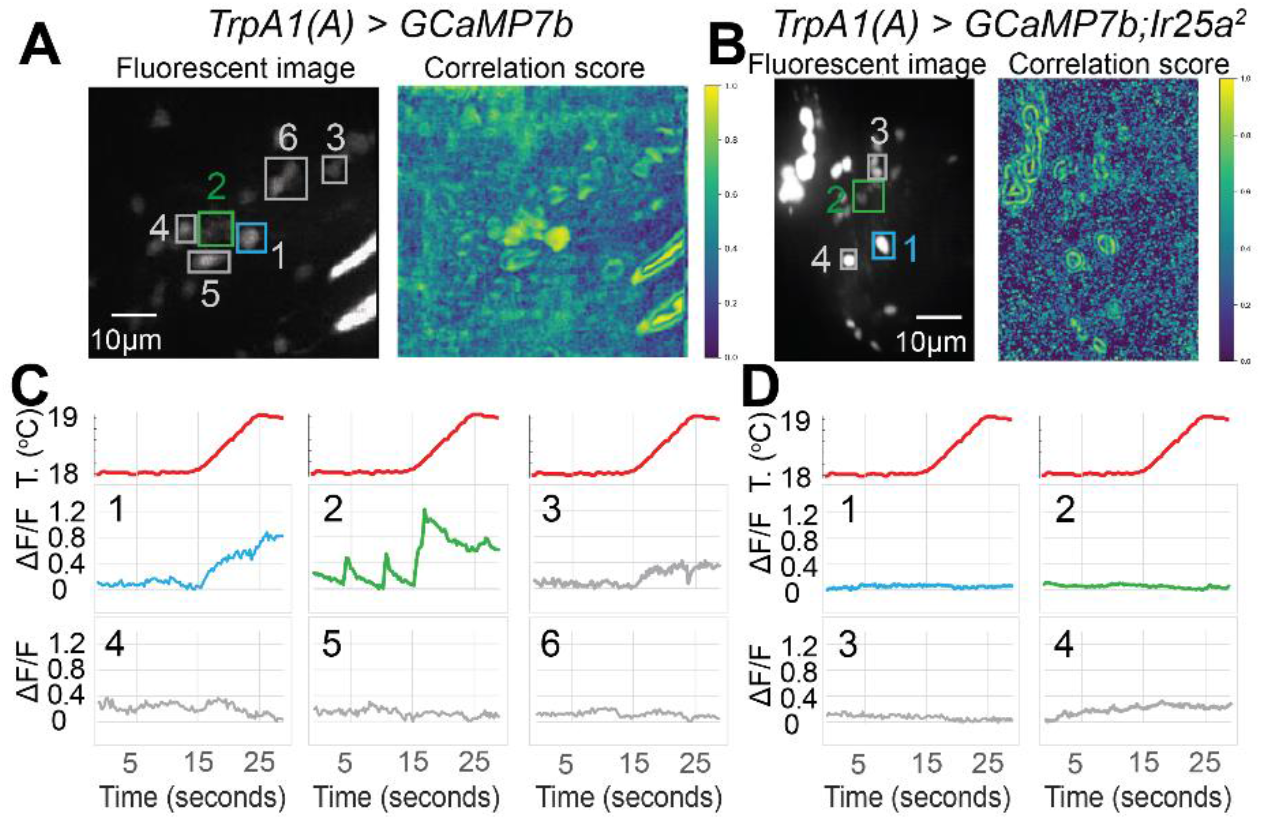
LPBTs do not respond to temperature increase in *Ir25a* mutants. **(A, B)** Examples of temperature responses in control **(A)** and *Ir25a*^2^ **(B)** preparations. Left, maximum fluorescence image with the regions of interest used for quantification outlined: LPBT cell bodies (blue), LPBT neurites (green) and other neurons in the field of view (grey). Right, local correlation image where each pixel is assigned the mean temporal correlation of its fluorescence trace with those of its immediate neighbors. Scale bars, 10 µm. **(C, D)** Fluorescence change (Δ*f/f*) in the regions outlined in **(A)** and **(B)** during a single 1 °C ramp from 18 °C (stage temperature in red). In control animals **(C)**, the neurites responded with transient increases during warming and the cell body with a slower, sustained rise, whereas other neurons in the same field did not respond. In *Ir25a*^2^ animals **(D)**, neither LPBT compartment responded. *n*=5 preparations for **A–D**

