## Supplementary Materials for "Gain Control of Peripheral Input by Core Brain Thermosensors Drives *Drosophila* Thermoregulation"

**The PDF file includes:**

Materials and Methods
Figs. S1 to S3
References (45-49)

### Materials and Methods

#### Fly strains

The standard laboratory strain Canton-S (CS) was used as the wild-type control. The strain $w^{1118}$, which provides the genetic background of the transgenic lines, was behaviorally indistinguishable from CS (Fig. S1). The following stocks were used: CS; $w^{1118}$ (BDSC 5905); *TrpA1*$^{1}$ (BDSC 36342); *TrpA1*$^{ins}$ from P.Garrity; *TrpA1(A)* (*TrpA1-A-KI*), *TrpA1(B)* (*TrpA1-B-KI*), *TrpA1(C)* (*TrpA1-C-KI*), *TrpA1(D)* (*TrpA1-D-KI*), *TrpA1-AD-KO*, *TrpA1-BC-KO*, *TrpA1-E-KO*, *TrpA1(WT)-GAL4* (*TrpA1-WT-KI-T2A-GAL4*), *TrpA1(A)-GAL4* (*TrpA1-A-KI-T2A-GAL4*), *TrpA1(C)-GAL4* (*TrpA1-C-KI-T2A-GAL4*) and *TrpA1(C)-LexA* (*TrpA1-C-KI-T2A-LexA*) from Y. Xiang; *TrpA1-AB-KO* (*TrpA1-AB-GAL4*) from C.Montell; *LexAop-FLP* from M. Rosbash; *Akh-GAL4* (BDSC 25684); *w;UAS-FRT-stop-FRT-CsChrimson::mVenus*; *w;UAS-FRT-stop-FRT-Kir2.1::GFP* from T.Shirangi(*1*); *UAS-GCaMP7b*; *Ir25a*$^{2}$; *UAS-syt-GCaMP6s*; *MB247-dsRed*; *R24E06-AD;R14E06-DBD* (SS01735), *R24E06-AD;VT045565-DBD* (SS01996), *R14E06-AD;R24E06-DBD* (SS01715) from M.Zlatic; *ChAT-GAL4;UAS-GFP* (BDSC 6793); *Gad1-EGFP* (BDSC 59304); *Vmat-EGFP* (BDSC 60263); *UAS-TrpA1-RNAi* (BDSC 36780). All mutant lines are homozygous viable and have been characterized previously in other contexts.

Stocks were maintained on standard medium (cornmeal, agar, yeast, corn syrup) at room temperature (approximately 23 °C). For experiments, adult males and virgin females of the relevant genotypes were placed in collection cages containing grape agar and yeast paste. Cages were kept at 23 °C and 60% relative humidity on a 12 h:12 h light:dark cycle and transferred to fresh medium every two days. Second-instar larvae, staged by spiracle morphology, were used for all experiments.

#### Temperature behavioral assay

Second-instar larvae were placed on the temperature stage inside a dark enclosure to exclude visual input. Ten to fifteen animals were recorded simultaneously for 20 min at 4 frames per second with a CCD camera; occasional collisions between animals were identified and excluded from the analysis. The stage consisted of a copper block with a 100 × 100 × 4 mm well filled with 2% agar containing 0.1% activated charcoal to increase contrast. The stage was heated and cooled by four Peltier devices backed by water blocks, through which antifreeze was circulated and cooled by a benchtop chiller (VWR). Peltier output was regulated by a proportional–integral–derivative controller using feedback from thermocouples mounted in the stage. Temperature was oscillated sinusoidally with a period of 60 s and an amplitude of ±1 °C about the median temperature.

Behavior was analyzed with the MAGAT Analyzer software package(*2*). For each larva in each frame, a midline spanning the length of the animal was constructed and segmented into 11 points; the positions of these points in consecutive frames were used to derive speed, direction of motion and head angle. Trajectories were then segmented into runs and turns in MATLAB (MathWorks) as described previously (*3*). Only trajectories lasting longer than 5 min were analyzed.

#### Behavioral metrics

Each frame was classified as turning or running. For each larva, the recording was folded onto the 60 s period of the temperature oscillation, and each time point within the cycle was assigned the category that occurred most often across cycles.

For each of the four phases of the cycle, the fraction of time turning was computed as the mean of the binary state over a ±3 s window centered on a fixed time point: warming, 17 s; peak, 34 s; cooling, 48 s; trough, 4 s, with t = 0 at the temperature minimum. The time point for the warming phase was chosen near the peak of the turning response of Canton-S larvae in $T_{mid}$ = 31 °C experiment and for cooling phase near the peak of the turning response in $T_{mid}$ = 23 °C. For the two static phases, the time points were chosen at the temperature maximum and minimum, with an additional 4 s offset applied to account for the latency of the behavioral response. The same time windows were applied across to all experiments. Change during heating was defined as the difference between the warming and trough values, and change during cooling as the difference between the cooling and peak values. For display, the per-larva time series were pooled and the fraction of larvae turning was computed at each time point, then smoothed with a five-point running mean. Smoothing was applied for display only and not before quantification.

#### Optogenetic experiments

Larvae carrying a GAL4 driver together with *UAS-CsChrimson* were reared in complete darkness in cages containing 2% agar and yeast paste supplemented with 0.5 mM all-*trans*-retinal. For experiments, animals were placed on the temperature stage held at a constant 18 °C and illuminated with modulated red light. A custom LED matrix ($\lambda$ = 650 nm) mounted above the stage provided uniform illumination with a maximum irradiance of 1.5 W $m^{-2}$, modulated sinusoidally with the same period as the thermal stimulus under custom hardware and software control (LabVIEW). Behavior was analyzed as for the temperature assay.

**Confocal microscopy and calcium imaging**

Imaging was performed on a spinning-disk confocal system (Nikon Instruments, Andor) with a 40×/1.4 NA oil-immersion objective. For calcium imaging, second-instar larvae were washed three times in distilled water and partially immobilized under a glass coverslip. Larvae were mounted on a copper stage with a steel plate, and temperature was regulated by a Peltier device with water cooling as in the behavioral assay. Temperature ramps of 1 °C were delivered at 0.1 °C $s^{-1}$, held for 20 seconds and returned at the same rate. Volumetric imaging was performed by stepping in z-axis with a piezo manipulator over a range of 40 µm with 20 steps and 100 Hz plane rate resulting in 5 volumes per second. Volumes were maximum-projected onto a single plane for quantification.

Motion correction and neuron tracking were performed with custom-built software in MATLAB, and regions of interest were defined from maximum projections. For Fig. 5E,F, motion correction, the correlation image and the regions of interest were generated with CaImAn(*4*) for display; these did not differ from the output of the custom code. Raw traces were corrected for bleaching by fitting a double-exponential decay to the fluorescence of non-responding neurons, background was subtracted using an area adjacent to the expressing tissue, and $\Delta F/F_0$ was computed with $F_0$ taken as the 5th percentile of the whole fluorescence trace for each region of interest.

**Immunohistochemistry**

For labeling LPBTs, brains of second-instar larvae carrying *TrpA1(C)-Gal4* and *UAS-mCD8::GFP* were dissected and fixed in electron-microscopy-grade 2% formaldehyde in phosphate-buffered saline (PBS) for 1 h. Tissues were washed three times for 1 h in PBS with 0.3% Triton X-100 (PBSTX), then blocked in PBSTX with 5% normal goat serum (NGS) for 30 min. Preparations were incubated with rotation at 4 °C over two nights in PBSTX containing 5% NGS, chicken anti-GFP (Aves Labs, 1:4000) and mouse anti-nc82 (Developmental Studies Hybridoma Bank, 1:40). Primary antibody staining was followed by three 20-min washes in PBSTX, after which tissues were incubated in goat anti-chicken Alexa Fluor 488 and goat anti-mouse Alexa Fluor 568 secondary antibodies (A-11039 and A-17004, ThermoFisher Scientific; 1:800 and 1:500 respectively) in PBSTX with 5% NGS at 4 °C over two nights before mounting in SlowFade (Invitrogen Molecular Probes). Samples were imaged on a Zeiss LSM 900 laser scanning confocal microscope. Images were analyzed using Arivis (Zeiss) and Fiji.

**Connectomics**

LPBTs were identified in a complete central nervous system from a 6-h-old Canton-S × $w^{1118}$ (BDSC 5905) larva acquired by serial-section transmission electron microscopy at a resolution of 3.8 × 3.8 × 50 nm(*5*). The wiring diagram was mapped using CATMAID(*6*). All neurons had been traced previously and synapse counts estimated. LPBTs were identified by morphological similarity and position among neurons downstream of the dorsal organ cool and warm cells, and the neurons so identified were used to select split-GAL4 drivers from an established collection(*7*). A threshold of five synapses was used to define synaptic partners, and the input fractions in Fig. 5C include all synapses identified in the connectome. We note that the connectome was reconstructed from a first-instar animal whereas all physiological and behavioral experiments used second-instar larvae.

**Statistics**

Each behavioral experiment was performed independently on at least three separate days and gave consistent results. The replication unit throughout is the individual larva, and sample sizes are given in the figures and figure legends. No statistical method was used to predetermine sample size. Investigators were not blinded to genotype, but all behavioral segmentation and all calcium-imaging source extraction were performed automatically by software without manual intervention.

Because the per-animal measures are bounded proportions with mass at the extremes, we did not assume normality. Confidence intervals on means were obtained by nonparametric bootstrap over animals (20,000 resamples, percentile method). Comparisons between genotypes at a given median or base temperature used two-sided Mann–Whitney *U* tests, with the rank-biserial correlation *r* reported as an effect size. Comparisons of paired measures within the same animal, and tests of a single measure against zero, used Wilcoxon signed-rank tests. Where a family of related comparisons was made across temperatures or across genotypes, *p* values were corrected within that family by the Holm–Bonferroni procedure, and corrected values are reported.

To test whether the dependence of a behavioral measure on median temperature differed between genotypes, we fitted ordinary least-squares models with a genotype by median-temperature interaction term, and in addition compared an additive model with a full interaction model by analysis of variance treating median temperature as a categorical factor, which makes no assumption of linearity. Both results are reported. The crossover temperature, defined as the median temperature at which turning during warming and during cooling are equal, was estimated from the zero crossing of a linear fit to the difference between them, with confidence intervals obtained by bootstrap stratified by temperature condition (5000 resamples).

In box plots, the center line shows the median and the box the interquartile range; whiskers extend to the most extreme observation within 1.5 times the interquartile range of the box edges, and all individual animals are shown as points. Throughout, * denotes $p < 0.05$, ** $p < 0.01$ and *** $p < 0.001$, and ns indicates $p \geq 0.05$. Analyses were performed in MATLAB (R2025b, MathWorks) and Python (3.9.12) using NumPy, SciPy, pandas and statsmodels.

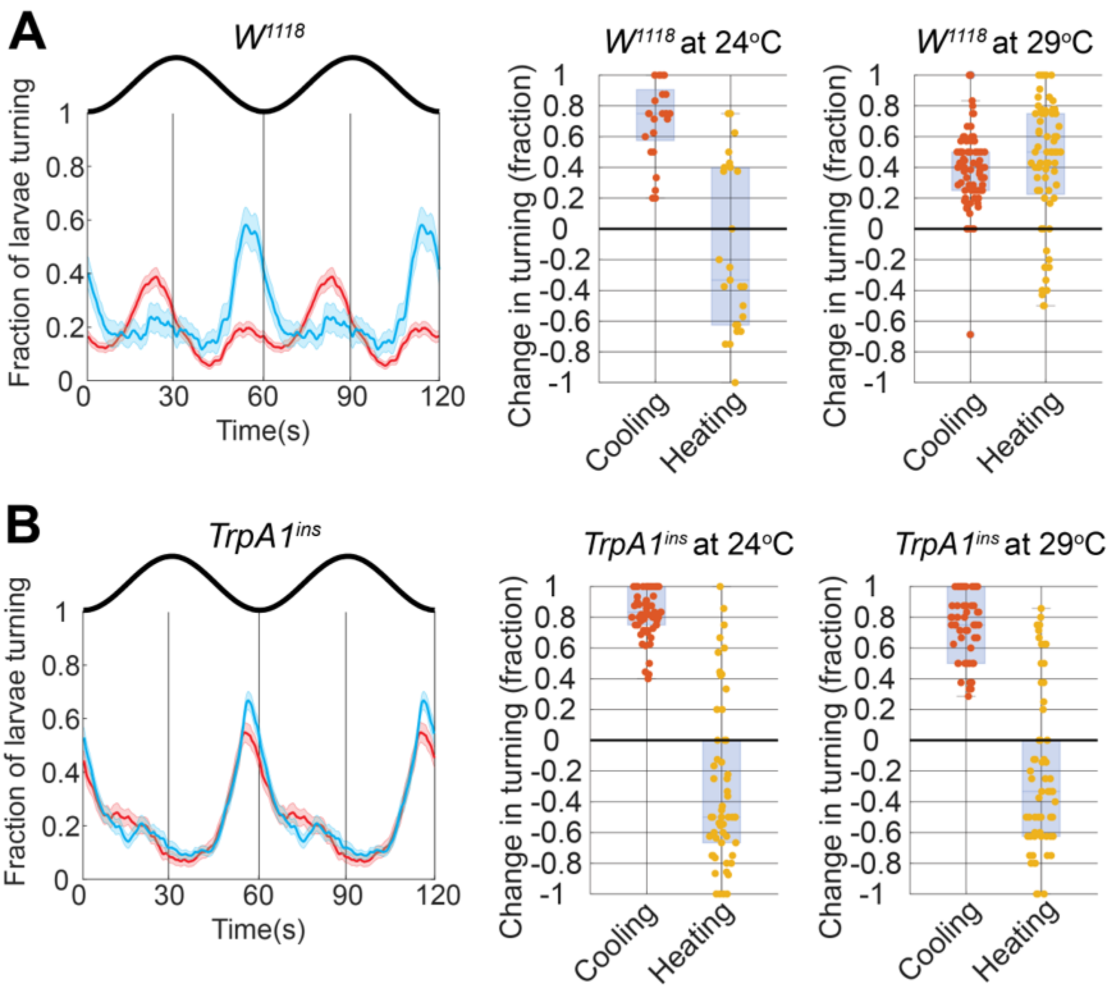


**Fig. S1. A second *TrpA1* allele reproduces the behavioral phenotype, while larvae mutant for the white gene do not. (A)** *TrpA1*$^{ins}$ larvae. **(B)** $w^{1118}$ mutant. Left, fraction of larvae turning across two oscillation cycles at $T_{\mathrm{mid}}$ = 24°C (blue) and 29 °C (red), with the stimulus waveform above; lines show the mean and bands the SEM. Center and right, change in turning during the cooling and warming intervals at each median temperature. Circles are individual larvae; boxes show the median and interquartile range. $w^{1118}$ larvae reversed the sign of the warming response between 24 °C and 29 °C, as Canton-S does, whereas *TrpA1*$^{ins}$ larvae turned during cooling at both median temperatures and never increased turning during warming. The phenotype therefore follows the *TrpA1* lesion rather than the genetic background. (n=36 for $w^{1118}$ at $T_{\mathrm{mid}}$ = 24°C, n = 84 for $w^{1118}$ at $T_{\mathrm{mid}}$ = 29°C, n=59 for *TrpA1*$^{ins}$ at $T_{\mathrm{mid}}$ = 24°C, n=62 for *TrpA1*$^{ins}$ at $T_{\mathrm{mid}}$ = 29°C)

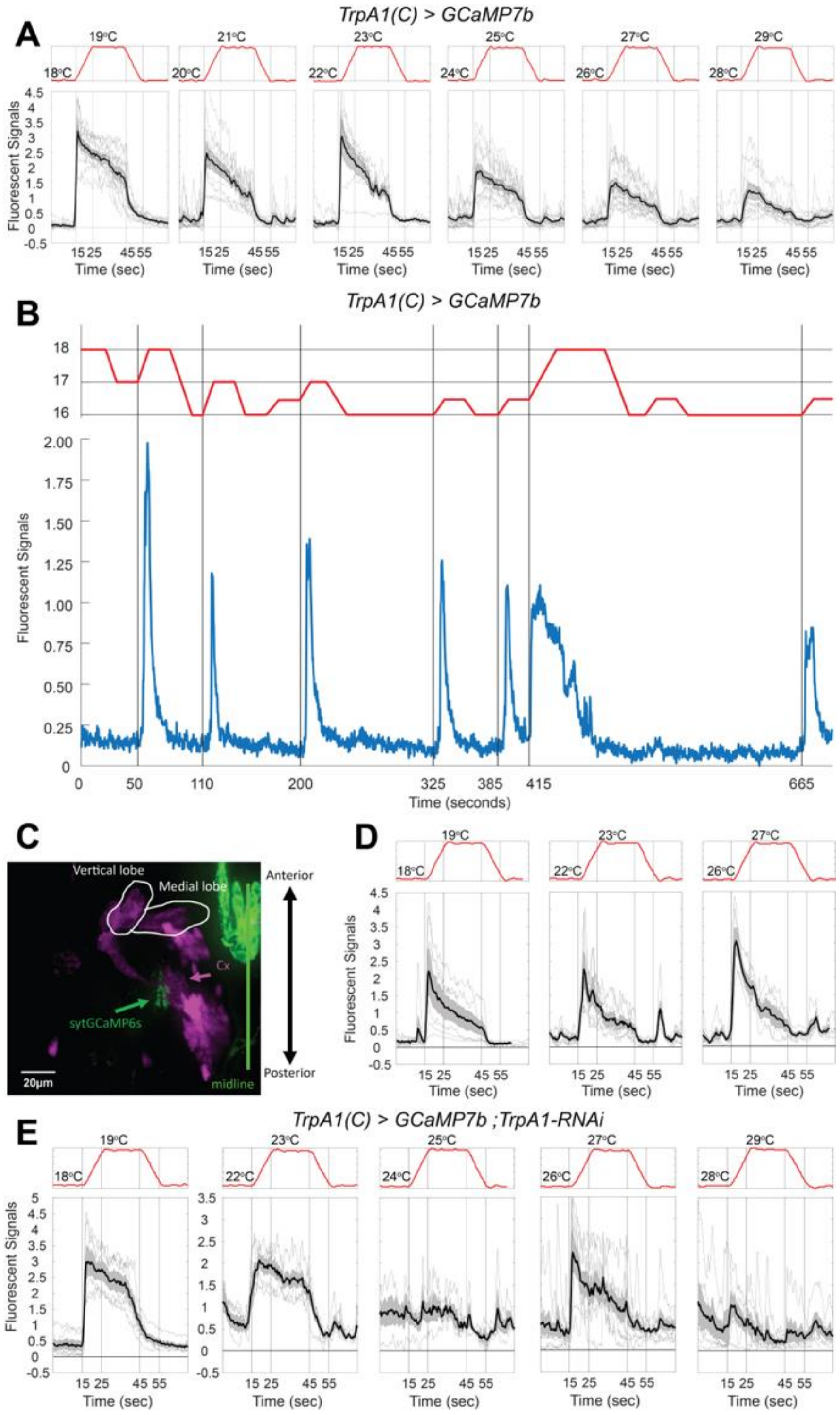


**Fig. S2. LPBT responses at various temperatures with *TrpA1(C)* driver, presynaptically expressed GCaMP and suppressed *TrpA1*. (A)** Responses to 1 °C ramps from six base temperatures in larvae expressing *GCaMP7b* under *TrpA1(C)-GAL4* control. The responses show reduced amplitude and decrease during temperature plateau unlike the signals from *TrpA1(A)-GAL4* driver. Due to the nature of the transgenic mutations, the *TrpA1(C) > GCaMP7b* animals only have one functional *TrpA1(A)* allele, while *TrpA1(A) > GCaMP7b* animals, used in the Fig. 4D, have both. **(B)** Stage temperature (red) and fluorescence (blue) in a single LPBT expressing GCaMP7b under *TrpA1(C)-GAL4* control, during a sequence of 1 °C and 2 °C steps between 16 °C and 18 °C. Each increase in temperature evoked a transient response, and no response followed a decrease. **(C)** Expression of synaptotagmin-GCaMP6s (green) driven by *TrpA1(C)-GAL4* in the lateral horn, with the mushroom body labeled in magenta; the mushroom body vertical and medial lobes and the calyx (Cx) are indicated as landmarks. Scale bar, 10 µm.

**(D)** Fluorescence responses in the LPBT terminal field expressing synaptotagmin-GCaMP6s to 1 °C ramps delivered from three base temperatures. **(E)** Responses to 1 °C ramps from five base temperatures in larvae expressing *TrpA1-RNAi* together with the calcium indicator under *TrpA1(C)-GAL4* control. Responses were retained at 18 °C and 22 °C but were reduced or absent from 24 °C upward, reproducing with cell-specific knockdown the temperature dependence seen in the *TrpA1*$^{1}$ null (Fig. 4). In **(A, D, E)**, thin lines are individual animals and the thick line and shaded band show the mean and SEM; stage temperature is shown in red above each panel.

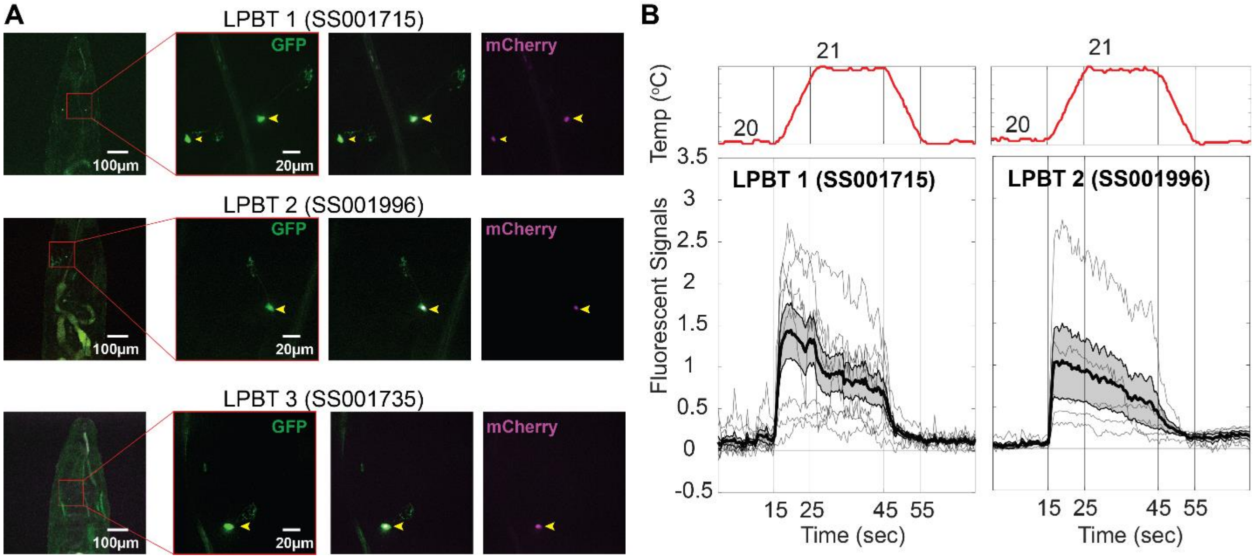


**Fig. S3. Split-GAL4 lines label LPBT neurons and reveal single-cell temperature responses.** (**A**) Three split-GAL4 combinations, selected on the basis of the connectome reconstruction, each label one or more of the three LPBT candidates: SS001715 (R14E06-AD;R24E06-DBD, LPBT 1), SS001996 (R24E06-AD;VT045565-DBD, LPBT 2) and SS001735 (R24E06-AD;R14E06-DBD, LPBT 3). For each line, left, overview of a whole-mount second-instar larva with the boxed region shown at higher magnification to the right; then membrane bound GFP (green), the merged image, and nuclear mCherry (magenta). Yellow arrowheads mark the labeled cell bodies. Scale bars, 100 µm (overviews) and 20 µm (insets). (**B**) Fluorescence responses of LPBT 1 and LPBT 2 to a 1 °C warming ramp from 20 °C (stage temperature in red above each panel). Thin lines are individual animals; the thick line and shaded band show the mean and SEM. Both neurons responded with a sustained increase during warming that decayed during cooling. No temperature-evoked responses were detected in LPBT 3. n = 5, 8 and 5 animals respectively.

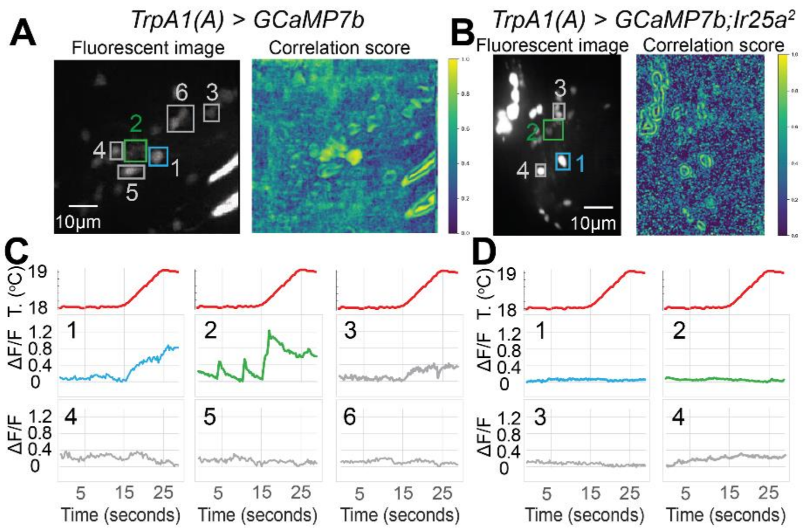


**Fig. S4. LPBTs do not respond to temperature increase in *Ir25a* mutants. (A, B)** Examples of temperature responses in control **(A)** and $Ir25a^2$ **(B)** preparations. Left, maximum fluorescence image with the regions of interest used for quantification outlined: LPBT cell bodies (blue), LPBT neurites (green) and other neurons in the field of view (grey). Right, local correlation image where each pixel is assigned the mean temporal correlation of its fluorescence trace with those of its immediate neighbors. Scale bars, 10 µm. **(C, D)** Fluorescence change ($\Delta f/f$) in the regions outlined in **(A)** and **(B)** during a single 1 °C ramp from 18 °C (stage temperature in red). In control animals **(C)**, the neurites responded with transient increases during warming and the cell body with a slower, sustained rise, whereas other neurons in the same field did not respond. In $Ir25a^2$ animals **(D)**, neither LPBT compartment responded. *n*=5 preparations for **A–D**
